# Protein family content uncovers lineage relationships and bacterial pathway maintenance mechanisms in DPANN archaea

**DOI:** 10.1101/2021.01.12.426361

**Authors:** Cindy J. Castelle, Raphaël Méheust, Alexander L. Jaffe, Kiley Seitz, Xianzhe Gong, Brett J. Baker, Jillian F. Banfield

**Affiliations:** Department of Earth and Planetary Science, University of California, Berkeley, CA, USA; Innovative Genomics Institute, University of California, Berkeley, CA, USA; Department of Plant and Microbial Biology, University of California, Berkeley, CA, USA; Department of Marine Science, University of Texas Austin, Port Aransas, TX, USA; Department of Environmental Science, Policy, and Management, University of California, Berkeley, CA, USA; Chan Zuckerberg Biohub, San Francisco, CA, USA; Institute of Marine Science and Technology, Shandong University, Qingdao, Shandong, China; LABGeM, Génomique Métabolique, Genoscope, Institut François Jacob, CEA, Evry, France

**Keywords:** DPANN, Archaea, Protein family, Bacterial genes, Phylogeny

## Abstract

DPANN are small-celled archaea that are generally predicted to be symbionts, and in some cases are known episymbionts of other archaea. As the monophyly of the DPANN remains uncertain, we hypothesized that proteome content could reveal relationships among DPANN lineages, constrain genetic overlap with bacteria, and illustrate how organisms with hybrid bacterial and archaeal protein sets might function. We tested this hypothesis using protein family content that was defined in part using 569 newly reconstructed genomes. Protein family content clearly separates DPANN from other archaea, paralleling the separation of Candidate Phyla Radiation (CPR) bacteria from all other bacteria. This separation is partly driven by hypothetical proteins, some of which may be symbiosis-related. Pacearchaeota with the most limited predicted metabolic capacities have Form II/III and III-like Rubisco, suggesting metabolisms based on scavenged nucleotides. Intriguingly, the Pacearchaeota and Woesearchaeota with the smallest genomes also tend to encode large extracellular murein-like lytic transglycosylase domain proteins that may bind and degrade components of bacterial cell walls, indicating that some might be episymbionts of bacteria. The pathway for biosynthesis of bacterial isoprenoids is widespread in Woesearchaeota genomes and is encoded in proximity to genes involved in bacterial fatty acids synthesis. Surprisingly, in some DPANN genomes we identified a pathway for synthesis of queuosine, an unusual nucleotide in tRNAs of bacteria. Other bacterial systems are predicted to be involved in protein refolding. For example, many DPANN have the complete bacterial DnaK-DnaJ-GrpE system and many Woesearchaeota and Pacearchaeota possess bacterial group I chaperones. Thus, many DPANN appear to have mechanisms to ensure efficient protein folding of both archaeal and laterally acquired bacterial proteins.

## Introduction

The first insights into archaeal cells of very small size and limited biosynthetic gene inventories predictive of symbiotic lifestyles were provided by researchers studying co-cultures of thermophilic *Ignicoccus* archaea and associated *Nanoarchaeum equitans* (Huber et al., 2002). Transmission electron microscope images established that these nanoarchaea are episymbionts (i.e., they attach to the surfaces of host *Ignicoccus* cells) (Jahn et al., 2008). Archaea with very small cell sizes and small genomes were discovered in acid mine drainage biofilms (Baker et al., 2006, 2010), apparently episymbiotically associated with Thermoplasmatales archaea (Comolli et al., 2009). Halophilic nanoarchaea were found in archaea-dominated planktonic communities (Narasingarao et al., 2012). Over time, a series of papers investigating the microbiology of sediments and aquatic environments greatly expanded the taxonomic diversity of nanoarchaea (Rinke et al., 2013a; Castelle et al., 2015a; Dombrowski et al., 2020). One group, groundwater-associated Huberarchaeota, are predicted to be symbionts of the archaeal Altiarchaeota (SM1) based on highly correlated abundance patterns (Probst et al., 2018). Recent work has further illuminated the biology, diversity and distribution of these organisms (Ortiz-Alvarez and Casamayor, 2016; Wurch et al., 2016; Krause et al., 2017;Dombrowski et al., 2019; Hamm et al., 2019), now often referred to as the DPANN (Rinke et al.,2013a). Genomic analyses were used to propose that lateral gene transfer, including from bacteria, has shaped the inventories of acidophilic nanoarchaea (Baker et al., 2010). This idea has been reinforced by more recent analyses of other DPANN archaea (Dombrowski et al.,2020; Jaffe et al., 2020).

The DPANN may comprise at least 12 different putative phylum-level lineages, but the phylogeny is still unresolved. For example, it is uncertain whether long branches within these groups are due to rapid evolution vs. undersampling. It also remains unclear which lineages are in vs. outside of the DPANN (Castelle et al., 2018) and whether lineages of archaea with small genomes form a monophyletic radiation (Aouad et al., 2018) analogous to the Candidate Phyla Radiation (CPR) of Bacteria or comprise multiple radiations. Additionally, while it has been suggested that Altiarchaeota may be part of the DPANN (Adam et al., 2017; Spang et al., 2017) there are currently no high quality genomes available for Altiarchaeota and the genomic sampling of this group is insufficient to resolve their placement. If Altiarchaeota are confirmed to be part of the DPANN, the metabolic characteristics of DPANN archaea would be expanded substantially (Probst et al., 2014).

Recently, researchers used HMM-based protein family-based analyses to provide insights regarding the relationships within and among major bacterial lineages, including those that comprise the bacterial CPR (Méheust et al., 2019). This approach has the advantage of being annotation-independent, thus can include consideration of proteins without functional predictions (Méheust et al., 2019). Here, we used a similar approach to investigate DPANN in the context of Domain Archaea. This analysis drew upon new genomes that we reconstructed for undersampled lineages, making use of metagenomic datasets from sediments and water. Our analyses of inventories of predicted DPANN proteins uncovered an intriguing number of bacteria-like sequences for functions typically associated with bacteria. The question of how proteome maintenance is achieved for organisms with a mixture of bacterial and archaeal systems was addressed in part by identification of bacterial-type protein refolding complexes in many DPANN genomes.

## Results

We collected 2,618 archaeal genomes from the NCBI genome database (**Supplemental Dataset - Table S1**) and augmented these by reconstructing 569 new DPANN draft genomes from low oxygen marine ecosystems, an aquifer adjacent to the Colorado River, Rifle, Colorado, and from groundwater collected at the Genasci dairy farm, Modesto, CA. The 3,197 genomes were clustered at ≥ 95% average nucleotide identity (ANI) to generate 1749 clusters. We removed genomes with <70% completeness or >10% contamination or if there was < 50% of the expected columns in the alignment of 14 concatenated ribosomal proteins (**see Materials and Methods**). We required that these proteins were co-encoded on a single scaffold to avoid contamination due to mis-binning of assembled genome fragments. Our analyses were performed on a final set that includes 390 DPANN genomes with an average completeness of 82% (**Supplementary Dataset - Table S1**).

We assessed the taxonomic distribution and affiliation of the 332 (348 if the genomes from (Parks et al., 2017) are included) newly reconstructed representative genomes based on the alignment of the syntenic block of 14 ribosomal proteins that is commonly used to infer genome-based phylogenies in previous studies (Castelle et al., 2015b) (**Figure 1, see Supplementary Figure 1 for more details**). These genomes were classified as DPANN because they cluster phylogenetically with known DPANN lineages or are most closely related to them (**Figure 1**). Addition of these new genome sequences increased the taxon sampling of the DPANN radiation and helped to resolve clades that were clustered together due to undersampling. The ‘Ca Mamarchaeota’ (Castelle and Banfield, 2018), now comprises 12 genome sequences from various ecosystem types and locations and is clearly monophyletic. We distinguished six new potential phylum-level DPANN lineages, the full definition of which will require further genomic sampling (**Figure 1**). The first is related to the Micrarchaeota phylum and is based on three genomes from groundwater collected at two different locations (Genasci and Rifle, USA). Two others, based on one and three genomes respectively, are deep branching with the Nanohaloarchaeota and were sampled from deep-subsurface marine sediment ecosystems. A fourth clade clustering at the root of the Nanoarchaeota and Parvarchaeota phyla includes seven genomes from marine and CO_2_-saturated groundwater (Crystal Geyser, Utah, USA). For this apparently phylum-level lineage we proposed the name of Candidatus “Kleinarchaeota”, named for the late Dr. Daniel Klein, a scientist who worked at the Joint Bioenergy Institute (**Figure 1**). Fifty new genome sequences further resolved two new clades within the Woesearchaeota phylum. The first clade corresponds to the Woesarchaeota-like II whereas the second clade groups together the Woesearchaeota and Woesarchaeota-like I groups.

**Figure 1.**
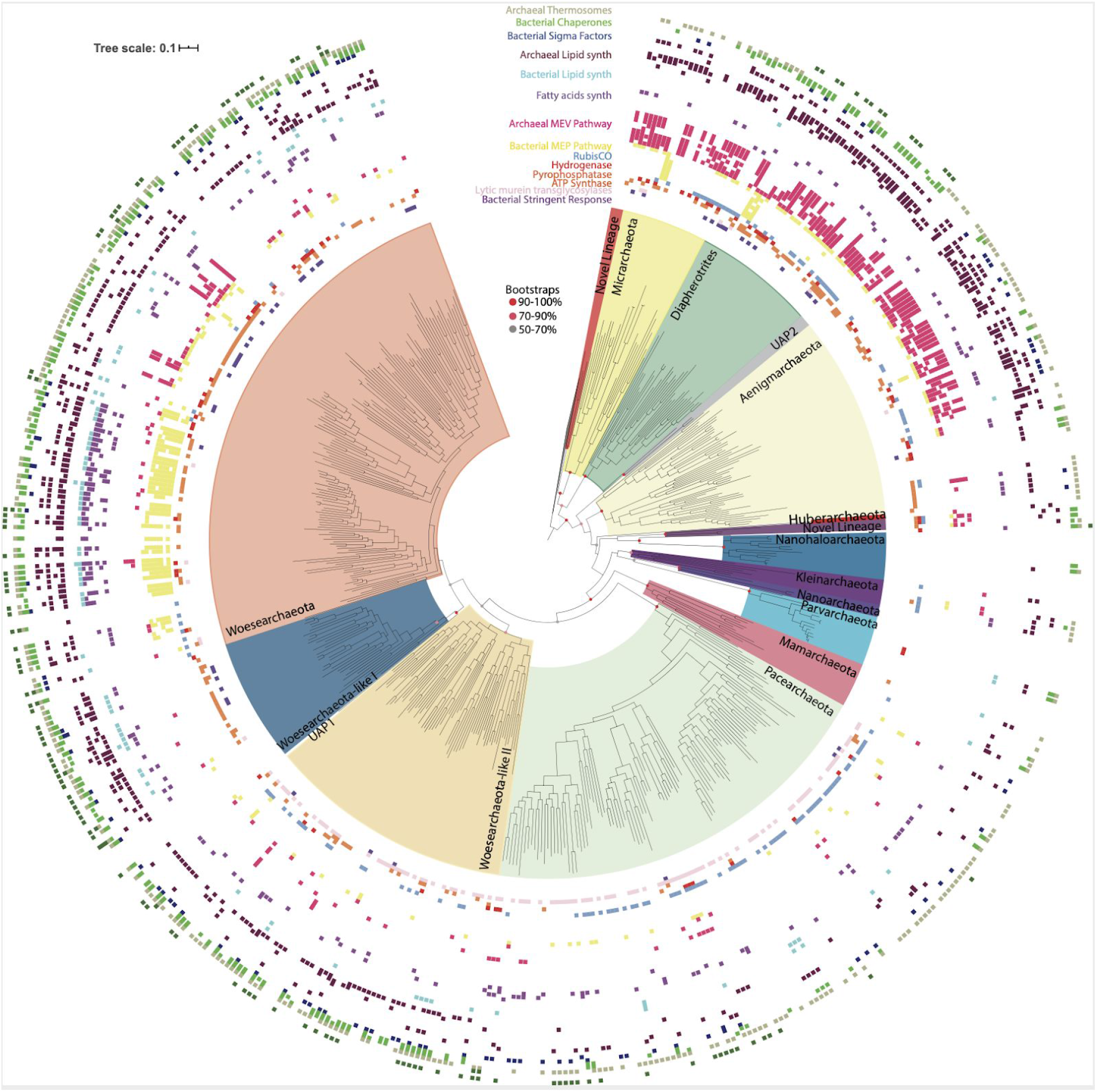
Taxonomic assessment and distribution of the newly DPANN genomes with metabolic similarities and distribution among DPANN lineages. Maximum-likelihood phylogeny of the DPANN lineages using Altiarchaeota as an outgroup and based on a 14-ribosomal-protein concatenated alignment (3677 amino acids, LG+R10 model). The presence/absence of a subset of targeted metabolic traits is indicated as concentric rings. The scale bar indicates the mean number of substitutions per site. A fully annotated tree with all included lineages, more metabolic features as well as the bootstrap values are available in **Supplementary Figure 1**.

## Protein clustering separates DPANN from other Archaea

We conducted an analysis of protein families, making use of a newly available set of 10,866 protein families for archaea (Méheust et al.). Analogous to the Candidate Phyla Radiation (CPR) bacteria, which appear to be the bacterial counterpart to the DPANN based on metabolic limitations, DPANN archaea separate from all other archaea based on their protein family contents (**Figure 2**) (Méheust et al., 2019). This is probably in part due to patterns of gene loss and their consistently minimal metabolic platforms.

**Figure 2.**
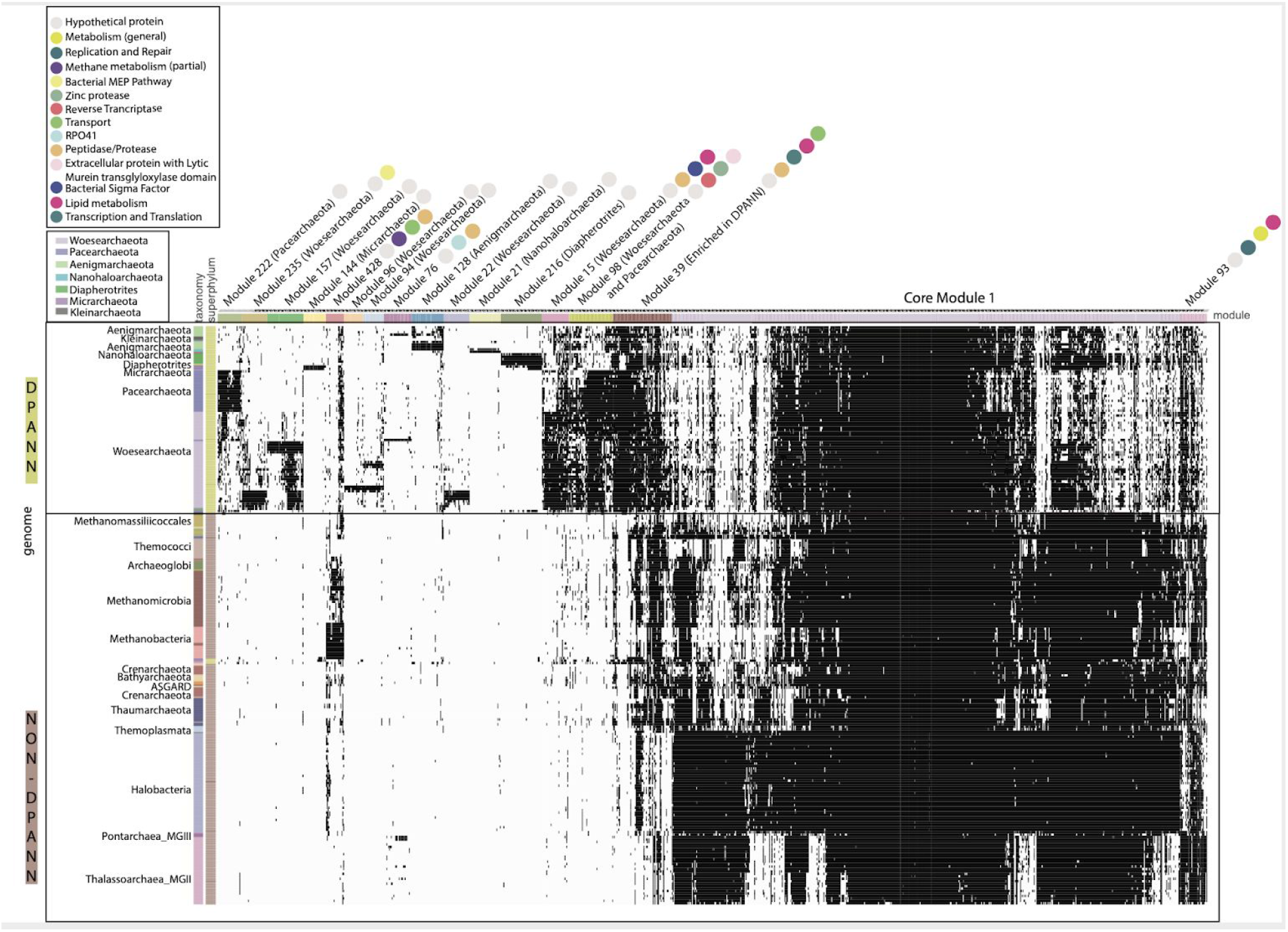
Distribution of 17 modules (504 Protein families) in DPANN genomes. The distribution of 504 protein families (columns) in representative genomes (rows) from Archaea. Data are clustered based on the presence (black) and absence (white) profiles (Jaccard distance, complete linkage). Pathway or function for each protein family is indicated by a colored circle.

The metabolic platforms differ from one DPANN lineage to another. Each module, a block of co-occurring protein families containing at least 20 families, was assigned a taxonomic distribution based on the taxonomy of the genomes with the highest number of families (**see Materials and Methods and Supplementary Dataset - Table S2**). Overall, 17 modules occurred in at least one DPANN genome (1,143 protein families), 14 modules are more common in DPANN compared to non-DPANN genomes (504 families), and 11 modules are unique to a specific DPANN lineage (**Table 1**) (**Figure 2**). Most of the protein families from those 14 modules have very poor or no functional annotations (**Supplementary Dataset - Table S3**). Importantly, we identified no modules that are common to all DPANN lineages but absent from other Archaea, although the module 39 is widespread in DPANN and only present in a few non-DPANN genomes (**Figure 2 and Supplementary Dataset - Table S2**).

**Table 1.** List of the twelve modules that are widely distributed within the genomes of DPANN archaea.

| Modules | Lineage(s) | # Families | SignalP (%) | TMHMM (%) | Hypothetical families (%) | Hits to Bacteria (%) |
| --- | --- | --- | --- | --- | --- | --- |
| 15,22,94,96,157,235 | Woesearchaeota | 178 | 19 | 48 | 93 | 15 |
| 98 | Woesearchaeota + Pacearchaeota | 51 | 24 | 53 | 90 | 39 |
| 216 | Diapherotrites | 48 | 25 | 73 | 98 | 2 |
| 128 | Aenigmarchaeota | 37 | 35 | 65 | 100 | 8 |
| 21 | Nanohaloarchaeota | 36 | 17 | 44 | 97 | 6 |
| 222 | Pacearchaeota | 28 | 36 | 79 | 100 | 18 |
| 144 | Micrarchaeota | 26 | 19 | 69 | 96 | 4 |

Interestingly, one module (Module 98) is widespread within Pacearchaeota and Woesearchaeota and unique to these groups. While most of the protein families in this module have unknown function, one protein family including proteins up to 3,000 aa in length (fam00969) contains a murein-like lytic transglycosylase domain potentially involved in cell defense and/or cell-cell interaction. This was previously noted in just three DPANN genomes (Rinke et al., 2013b; Castelle et al., 2015b). Murein transglycosylases are near-ubiquitous in Bacteria (but normally absent in Archaea) and specifically bind and degrade murein (peptidoglycan) strands, the main components of the bacterial cell walls. Thus, these large proteins might be a novel class of extracellular enzymes specific to Woesearchaeota and Pacearchaeota and involved in attachment to bacterial cell walls, potentially including those of their hosts.

Overall, the vast majority of the DPANN lineage-specific protein families lack function predictions. For instance, the Pacearchaeota and Aenigmarchaeota each had lineage-specific modules (Modules 222 and 128 respectively) completely lacking functional predictions (**Table 1**). Despite the lack of functional annotations, many of the DPANN lineage-specific protein families are predicted to be either membrane-bound or extracellular (**Table 1** and **Supplementary Dataset - Table S3**).

## Bacterial systems in DPANN archaea

To determine the extent to which inter-domain lateral transfer has impacted metabolic capacity in DPANN archaea, we computed a metric (“breadth”) that describes the incidence of each DPANN protein family across domain Bacteria (**Figure 3**) (Materials and Methods). Most of those with the highest incidence across Bacteria (as measured by “breadth”) are involved in core biological functions that are common to both bacteria and archaea (module 1) (**Supplementary dataset - Table S3, Material and Methods**). On the other hand, we also observed a number of smaller archaeal protein families that were relatively common in Bacteria (**Figure 3**) and these bacterial-like sequences were more prominent in DPANN compared to other archaea. We reasoned that these uncommon archaeal protein families could either reflect contamination introduced by misbinning or inter-domain lateral gene transfer from Bacteria to Archaea. Thus, we retained them for further analysis. Interestingly, only a few DPANN protein families with unknown or uncharacterized function(s) were detected in bacteria. This could suggest that the specificity of the majority of families of hypothetical proteins are unique to DPANN Archaea; however, we cannot fully exclude the possibility that homology was not detected due to high divergence.

**Figure 3.**
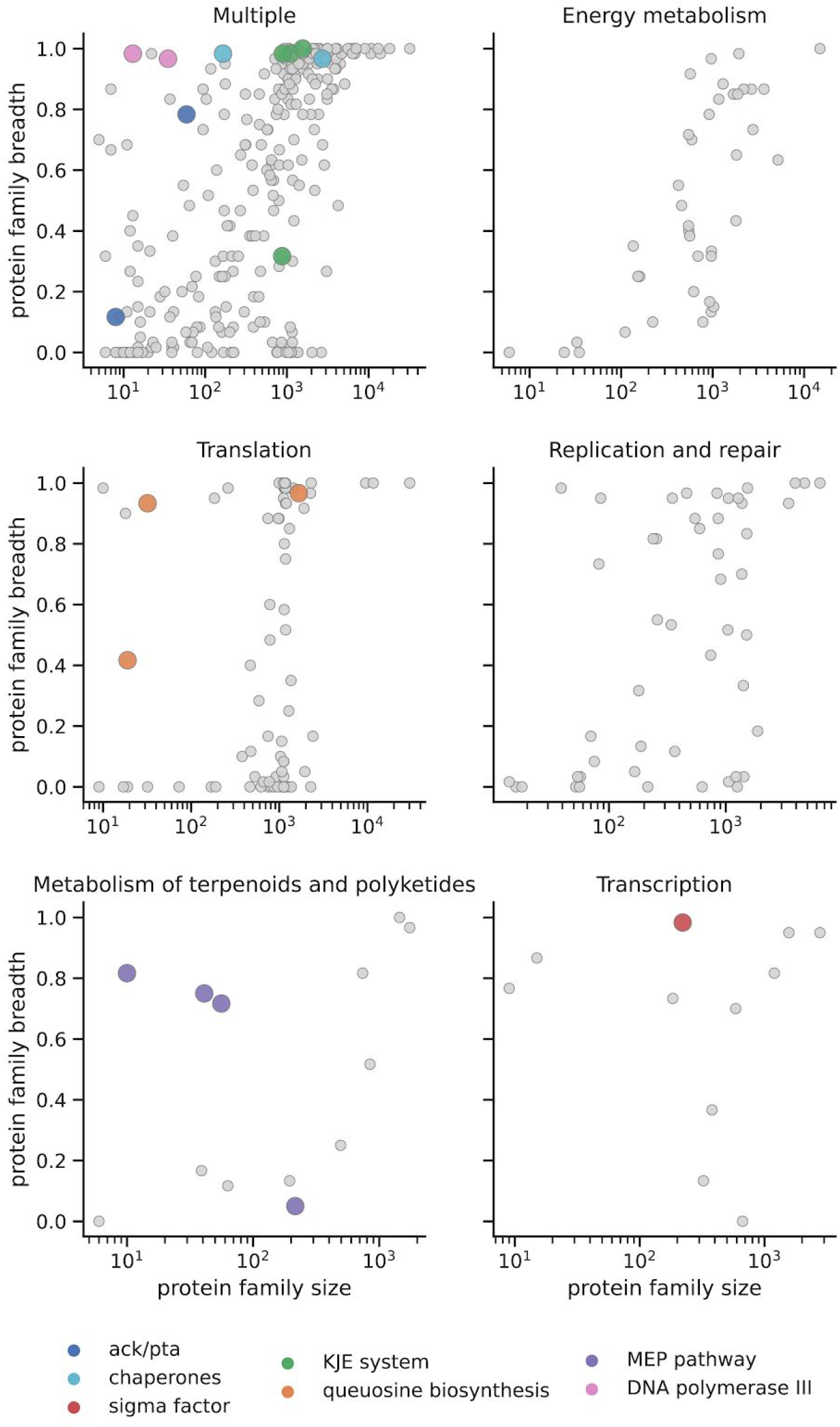
Bacterial protein families found in DPANN genomes. In the top left plot, “multiple” refers to protein families that are found in more than one metabolic pathway. Colored dots indicate those instances in DPANN. For example, translation genes in the DPANN (pink dots) are genes involved in queosine biosynthesis (see key).

We then further examined small archaeal protein families with a strong signal of bacterial origin (“breadth”), focusing on those with confident functional annotations. Some bacterial systems involved in transcription, protein folding, and cell membrane production are encoded in many DPANN archaeal genomes. For example, in the newly reconstructed genomes we uncovered genes for sigma factors classified as σ-70 (fam00359, **Supplementary dataset - Table S3**). These are particularly prevalent in Woesearchaeota (Module 15). Their presence due to mis-binning was ruled out based on the phylogeny of co-encoded and taxonomically informative proteins. We did not identify any possible anti-sigma factors, which are essential for controlling and regulating sigma factors in Bacteria. Thus, if involved in transcription regulation (along with the expected archaeal transcription factors TBP, TFB, TFE that are also found in DPANN), the mechanism controlling them remains unclear.

Other bacterial systems identified in DPANN genomes are the GroEL and Hsp60 group I chaperones involved in protein folding. Group II chaperones consist of the archaeal thermosomes and related eukaryotic cytosolic variants (CCT or TRiC) (Lopez et al., 2015) and generally do not coexist with Group I chaperones in the same cell. Analysis of DPANN genomes revealed that many Woesearchaeota and Pacearchaeota possess both archaeal thermosomes and GroEL/ES (fam00693 and fam02048, respectively). A similar observation was made for some species of mesophilic archaea (methanogens of the order of Methanosarcina) (Figueiredo et al., 2004). The presence of both systems may ensure efficient protein folding of both traditionally archaeal and laterally acquired bacterial proteins.

Another well-known chaperone system in bacteria is the DnaK-DnaJ-GrpE (KJE) system, which funnels its clients toward the native state or ushers misfolded proteins into degradation. Within Domain Archaea, the KJE system only has been reported in a few archaeal species and was likely acquired by horizontal gene transfer (Gribaldo et al., 1999; Petitjean et al., 2012). The KJE system has been considered an essential building block for a minimal bacterial genome and has been reported to be mostly absent from highly reduced endosymbionts or obligate bacterial pathogens (McCutcheon and Moran, 2011). Surprisingly, we find that many DPANN archaea have the complete bacterial KJE system (Module 1: DnaK:fam00458; DnaJ: fam00163 and fam00567; GrpE: fam01146, **Supplementary dataset - Table S3**)

Previously, we reported the distribution of archaeal and bacterial isoprenoid synthesis pathways (the archaeal mevalonate (MVA) and the bacterial 2-C-methyl-D-erythritol 4-phosphate/1-deoxy-D-xylulose 5-phosphate (MEP/DOXP) pathways) in many DPANN genomes (Castelle and Banfield, 2018). Inclusion of hundreds of new DPANN genomes in this study allowed us to further evaluate the distribution of the MEP pathway. Interestingly, we found this pathway is widespread in a monophyletic clade composed of 52 Woesearchaeota genomes (**Figure 1**) that were reconstructed from a wide range of terrestrial and marine ecosystems. The genes involved in the biosynthesis of bacterial isoprenoids appear to be co-located in Woesearchaeota genomes (**Supplementary Dataset - Table S4**). In proximity to the MEP pathway genes we identified a gene encoding for an UbiA prenyltransferase (fam00115), which is involved in the synthesis of archaeal ether lipids, and transfers prenyl groups to hydrophobic ring structures such as quinones, hemes, chlorophylls, vitamin E or shikonin (Villanueva et al.,2014). Some of these archaea have genes involved in bacterial fatty acid synthesis, including the bacterial glycerol-3-Phosphate dehydrogenase, and in some genomes they are co-located with MEP pathway genes (G3P; fam00008; **Supplementary Dataset - Table S4**).

One of the most common predictions for the metabolic basis for growth of DPANN archaea is fermentation (Castelle et al., 2015b, 2018). For instance, many DPANN archaea are capable of producing acetate, primarily via a single enzyme, an acetyl-CoA synthetase that is the most common enzyme for this function found in Archaea. On the other hand, most Bacteria use a pathway involving acetate kinase (Ack) and phosphotransacetylase (Pta), which is the reversible reaction allowing Bacteria to oxidize or produce acetate. The only Archaea currently known to use the Ack/Pta pathway are members of the methanogenic Methanosarcina (Fournier and Gogarten, 2008). Here, we report that the Ack/Pta pathway occurs in many Pacearchaeota and Woesearchaeota (fam06325 and fam12605, respectively) (**Supplementary Dataset - Table S3**). Thus, these DPANN archaea have the capacity to both excrete and assimilate acetate, as occurs in some bacteria. This may provide an adaptive advantage under changing environmental conditions by allowing growth by fermentation of abundant nutrients and enhanced survival on acetate when nutrients are depleted.

Transfer RNA (tRNA) is structurally unique among nucleic acids in harboring an astonishing diversity of post-transcriptionally modified nucleosides. Two of the most radically modified nucleosides known to occur in tRNA are queuosine and archaeosine, both of which are characterized by a 7-deazaguanosine core structure (Itaya, 2003). In spite of the phylogenetic segregation observed for these nucleosides (queuosine is present in Eukarya and Bacteria, while archaeosine is present only in Archaea), their structural similarity suggested a common biosynthetic origin. Surprisingly, we identified the complete and/or partial bacterial queuosine biosynthesis pathway in 34 archaeal genomes (including 12 DPANN genomes). In DPANN, this queuosine synthesis pathway includes co-localized (**Figure 4 and Supplementary Dataset - Table S5**) QueA (S-adenosylmethionine:tRNA ribosyltransferase-isomerase; fam24423), QueH-like (epoxyqueuosine reductase; fam24901) and Tgt (queuine tRNA-ribosyltransferase; Fam00366).

**Figure 4.**
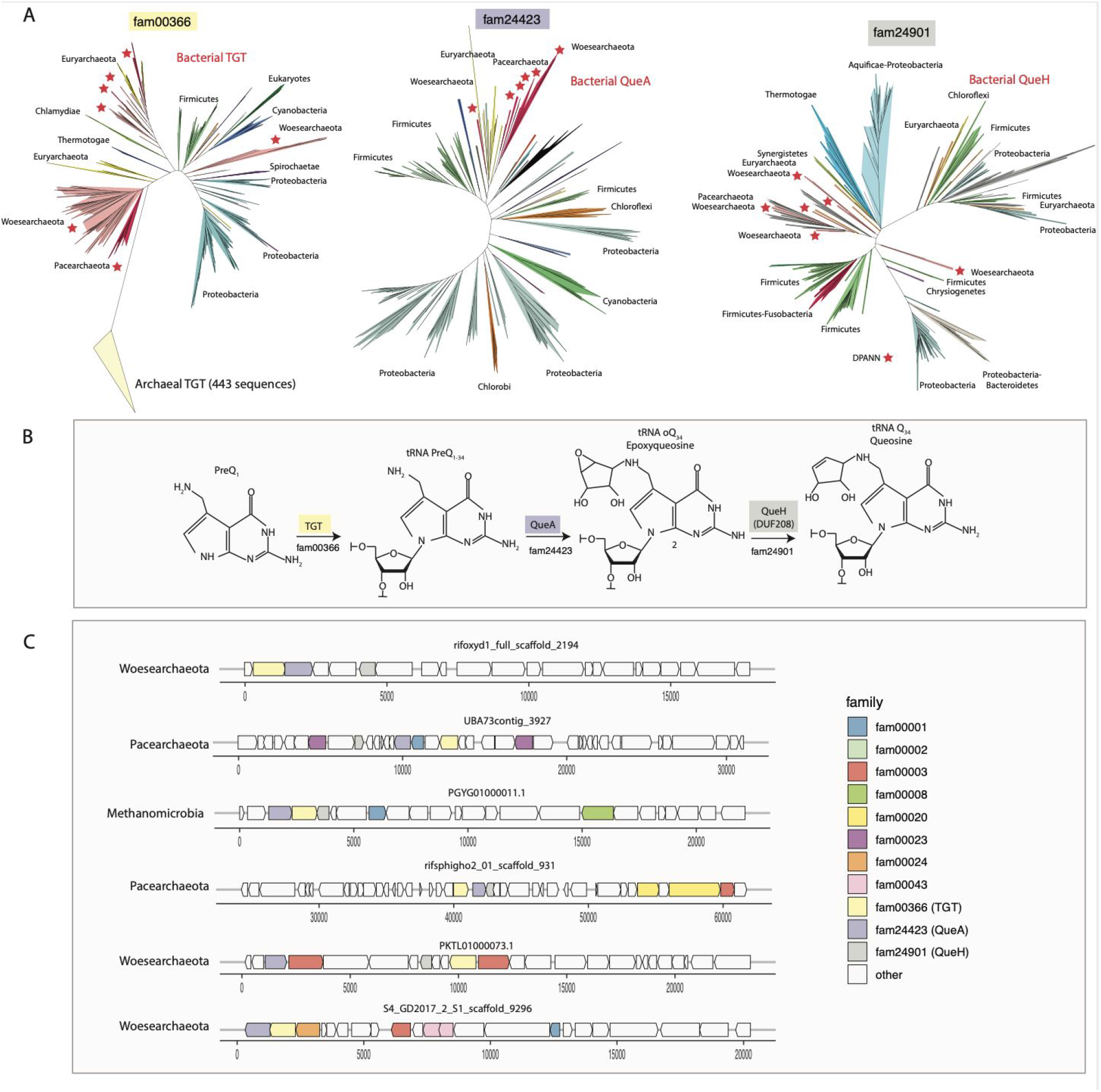
Bacterial queuosine pathway in DPANN genomes and some Euryarchaeota (Methanosarcinales). **A**, Phylogenetic affiliation and distribution of the identified QueA, QueH and TGT in archaea selected for this study. **B**, The three last steps of the queuosine biosynthesis pathway in Bacteria. **C**, Selected examples of genomic context and gene synteny around the genes QueA, QueH (DUF208) and TGT. QueA, S-adenosylmethionine:tRNA ribosyltransferase-isomerase; QueH (DUF208), epoxyqueuosine reductase; TGT, queuine tRNA-ribosyltransferase. A full gene synteny and genomic context of all the genes involved in the queuosine pathway in this study is available in **Supplementary Dataset - Table S5**.

Some DPANN genomes encode DNA polymerase III, a bacterial polymerase not typically found in Archaea. The DPANN sequences were verified as part of DPANN genomes based on the presence of archaeal ribosomal proteins and other phylogenetically informative genes in close genomic proximity. Placing these sequences in phylogenetic context reveals that some sequences from Aenigmarchaeota cluster with sequences from Chloroflexi, possibly reflecting lateral transfer from this group (**Figure 5**). Additionally, the Pacearchaeota sequences are highly divergent from bacterial sequences, yet the domain structure supports their classification as DNA polymerase III.

**Figure 5.**
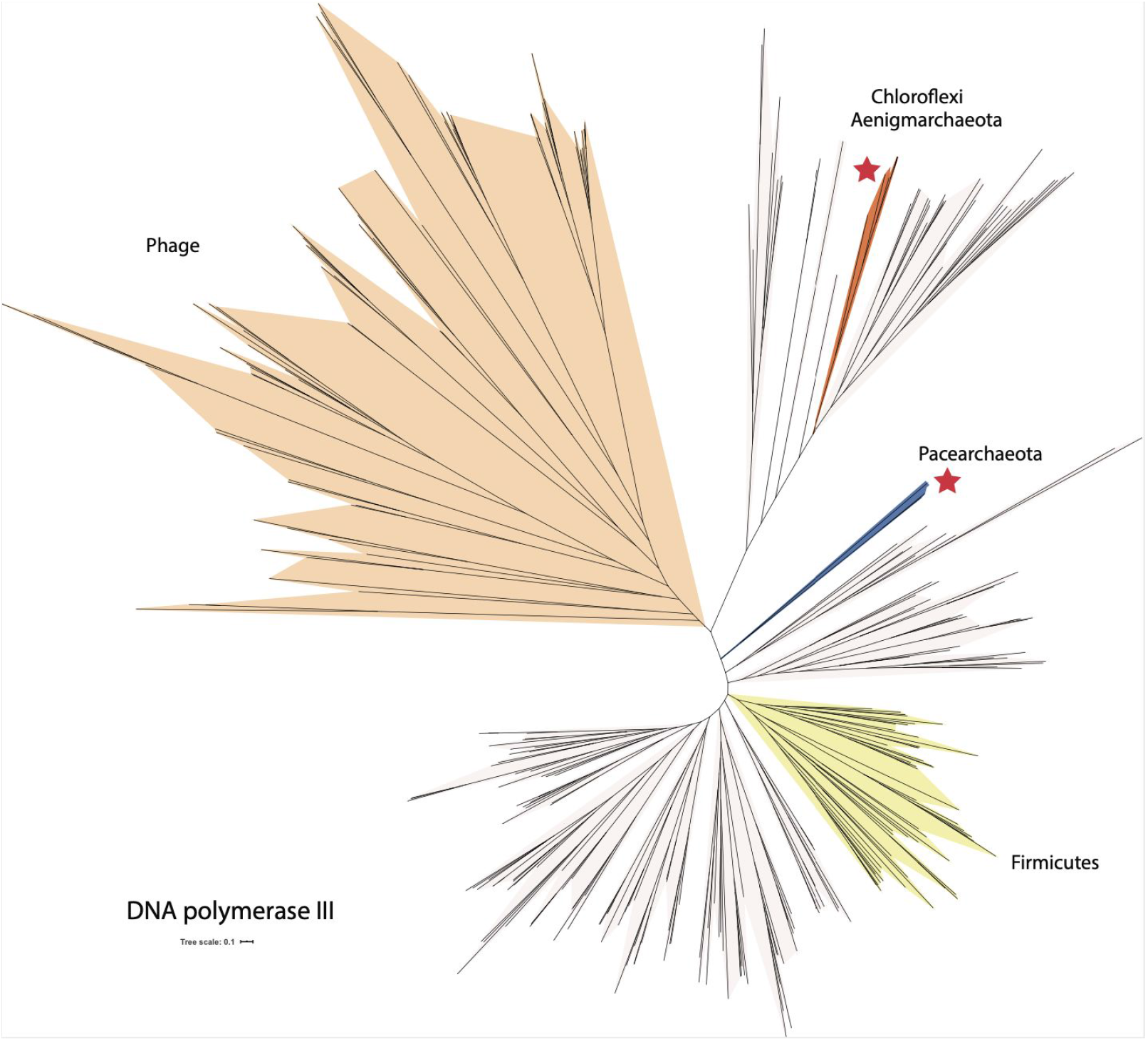
Phylogenetic tree of DNA polymerase III, a bacterial polymerase not typically found in Archaea. The tree includes sequences from phages and some bacteria (light cream shading). Asterisks indicate groups of sequences found in DPANN genomes. The scale bar indicates the mean number of substitutions per site.

## Conclusions

Much remains to be learned about DPANN archaea, and the question of whether they share a common ancestor remains unclear. We found no set of DPANN-specific protein families, as was observed for the CPR bacteria. However, some modules of protein families are shared by multiple DPANN distinct lineages. For example, Woesearchaeota and Pacearchaeota share distinct protein family modules, and although they are clearly separate phylogenetic lineages and have distinct metabolism capacities. Thus, these lineages, at least, could have shared a common ancestor from which they inherited a set of lineage specific proteins.

Based on the protein family analysis, DPANN archaea are distinct from other archaea. This may be in part due to patterns of gene loss as well as lateral gene transfer. Similar patterns of gene loss on the path to symbioses could have arisen via convergent evolution or could indicate that phylogenetically distinct lineages have shared ancestry. Our findings related to lateral gene transfer extend those of a very recent report that suggested the phylogenetic analyses of DPANN archaea may be confounded by this process (Dombrowski et al., 2020). However, our results suggest that sets of proteins, including those with known and unknown functions, were established at the origins of the major lineages and have been largely preserved within each phylum.

The question of why DPANN archaea have acquired bacterial proteins is intriguing, and may indicate close physical associations between DPANN and bacteria. It remains to be seen whether, in some cases, DPANN have bacterial hosts. The presence in DPANN proteomes of typically bacterial systems, including proteins potentially involved in genome regulation and protein refolding, may explain in part how these hybrid protein inventories are maintained. The findings hint at the existence of mechanisms that enable lifestyles based on close associations that involve organisms from the bacterial and archaeal domains. Furthermore, the presence of bacterial systems even in reduced DPANN genomes is interesting, considering that large-scale bacterial gene import has been suggested to underlie the origin of major archaeal lineages (Nelson-Sathi et al., 2015)).

## Materials and Methods

### Genome collection

569 unpublished genomes were added to the 2,618 genomes of Archaea downloaded from the NCBI genome database in September 2018 (**Supplementary Dataset - Table S1**).

One hundred thirty-two genomes were obtained from metagenomes of sediment samples. Sediment samples were collected from the Guaymas Basin (27°N0.388, 111°W24.560, Gulf of California, Mexico) during three cruises at a depth of approximately 2,000 m below the water surface. Sediment cores were collected during two Alvin dives, 4486 and 4573 in 2008 and 2009. Sites referred to as “Megamat” (genomes starting with “Meg”) and “Aceto Balsamico” (genomes starting with “AB” in name), Core sections between 0-18 cm from 4486 and from 0-33 cm 4573 and were processed for these analyses. Intact sediment cores were subsampled under N2 gas, and immediately frozen at −80 °C on board. The background of sampling sites was described previously (Teske et al., 2016). Samples were processed for DNA isolation from using the MoBio PowerMax soil kit (Qiagen) following the manufacturer’s protocol. Illumina library preparation and sequencing were performed using Hiseq 4000 at Michigan State University. Paired-end reads were interleaved using interleav_fasta.py (https://github.com/jorvis/biocode/blob/master/fasta/interleave_fasta.py) and the interleaved sequences were trimmed using Sickle (https://github.com/najoshi/sickle) with the default settings. Metagenomic reads from each subsample were individually assembled using IDBA-UD with the following parameters: --pre_correction --mink 65 --maxk 115 --step 10 --seed_kmer 55 (Peng et al., 2012). Metagenomic binning was performed on contigs with a minimum length of 2,000 bp in individual assemblies using the binning tools MetaBAT (Kang et al., 2015) and CONCOCT (Alneberg et al., 2014), and resulting bins were combined with using DAS Tool (Sieber et al.). CheckM lineage_wf (v1.0.5) (Parks et al., 2015) was used to estimate the percentage of completeness and contamination of bins. Genomes with more than 50% completeness and 10% contamination were manually optimized based on GC content, sequence depth and coverage using mmgenome (Karst et al.).

One hundred eighty-eight genomes were obtained from groundwater samples from Genasci Dairy Farm, located in Modesto, California (CA), USA (He et al.). Over 400 L of groundwater were filtered from monitoring well 5 on Genasci Dairy Farm over a period ranging from March 2017 to June 2018. DNA was extracted from all filters using Qiagen DNeasy PowerMax Soil kits and ~10 Gbp of 150 bp, paired end Illumina reads were obtained for each filter. Assembly was performed using MEGAHIT (Li et al., 2015) with default parameters, and the scaffolding function from assembler IDBA-UD was used to scaffold contigs. Scaffolds were binned on the basis of GC content, coverage, presence of ribosomal proteins, presence of single copy genes, tetranucleotide frequency, and patterns of coverage across samples. Bins were obtained using manual binning on ggKbase (Wrighton et al., 2012), Maxbin2 (Wu et al., 2016), CONCOCT (Alneberg et al., 2014), Abawaca1, and Abawaca2 (https://github.com/CK7/abawaca), with DAS Tool (Sieber et al.) used to choose the best set of bins from all programs. All bins were manually checked to remove incorrectly assigned scaffolds using ggKbase.

Additionally, 168 genomes were obtained from an aquifer adjacent to the Colorado River near the town of Rifle, Colorado, USA, at the Rifle Integrated Field Research Challenge (IFRC) site (Anantharaman et al., 2016). Sediment samples were collected from the ‘RBG’ field experiment carried out in 2007. Groundwater samples were collected from three different field experiments. All groundwater samples were collected from 5 m below the ground surface by serial filtration onto 1.2, 0.2 and 0.1 μm filters (Supor disc filters; Pall Corporation, Port Washington, NY, USA). DNA was extracted from all frozen filters using the PowerSoil DNA Isolation kit (MoBio Laboratories Inc., Carlsbad, CA, USA) and 150 bp paired end Illumina reads with a targeted insert size of 500 bp were obtained for each filter. Assemblies were performed using IDBA-UD (Peng et al., 2012) with the following parameters: --mink 40, --maxk 100, --step 20, --min_contig 500. All resulting scaffolds were clustered into genome bins using multiple algorithms. First, scaffolds were binned on the basis of % GC content, differential coverage abundance patterns across all samples using Abawaca1, and taxonomic affiliation. Scaffolds that did not associate with any cluster using this method were binned based on tetranucleotide frequency using Emergent Self-Organizing Maps (ESOM) (Dick et al., 2009). All genomic bins were manually inspected within ggKbase.

Fifty genomes were obtained from the Crystal Geyser system in Utah, USA (Probst et al., 2018). Microbial size filtration from Crystal Geyser fluids was performed using two different sampling systems. One system involved sequential filtration of aquifer fluids on 3.0-μm, 0.8-μm, 0.2-μm, and 0.1-μm filters (polyethersulfone, Pall 561 Corporation, NY, USA). The second system was designed to filter high volumes of water sequentially onto 2.5-μm, 0.65-μm, 0.2-μm and 0.1-μm filters (ZTECG, Graver Technologies, Glasgow, USA). Metagenomic DNA was extracted from the filters using MoBio PowerMax soil kit. DNA was subjected to 150 bp paired end illumina HiSeq sequencing at the Joint Genome Institute. Assembly of high-quality reads was performed using IDBA_UD with standard parameters and genes of assembled scaffolds (>1kb). Genome bins were obtained using different binning algorithms: Semi-automated tetranucleotide-frequency based emergent self-organizing maps (ESOMs), differential coverage ESOMs, Abawaca1, MetaBAT and Maxbin2. Best genomes from each sample were selected using DASTool. All bins were manually checked to remove incorrectly assigned scaffolds using ggKbase.

Finally, forty one genomes were obtained from the Uncultivated Bacteria and Archaea project (Parks et al., 2017)but were manually curated using ggKbase.

### Genome completeness assessment and de-replication

Genome completeness and contamination were estimated based on the presence of single-copy genes (SCGs). Genome completeness was estimated using 38 SCGs (Anantharaman et al., 2016). DPANN genomes with more than 22 SCGs and less than 4 duplicated copies of the SCGs were considered as draft-quality genomes. Genomes were de-replicated using dRep (Olm et al., 2017) (version v2.0.5 with ANI > 95%). The most complete and less contaminated genome per cluster was used in downstream analyses.

### Concatenated 14 ribosomal proteins phylogeny

A maximum-likelihood tree was calculated based on the concatenation of 14 ribosomal proteins (L2, L3, L4, L5, L6, L14, L15, L18, L22, L24, S3, S8, S17, and S19). Homologous protein sequences were aligned using MAFFT (version 7.390) (--auto option) (Katoh and Standley,2016). The protein alignments were concatenated and manually refined, with a final alignment of 551 genomes and 3,677 positions. Phylogenetic tree was inferred using RAxML (version 8.2.10) (Stamatakis, 2014) (as implemented on the CIPRES web server (Miller et al., 2010)), under the LG+R10 model of evolution, and with the number of bootstraps automatically determined.

### Module definition and taxonomic assignment

Looking at the distribution of the protein families across the genomes, a clear modular organization emerged. Modules of families were defined using a cutoff of 0.95 on the dendrogram tree of the families. The dendrogram tree was obtained from a hierarchical clustering using the Jaccard distance that was calculated based on profiles of protein family presence/absence. The corresponding clusters define the modules.

A phyla distribution was assigned to each module using the method of (Méheust et al., 2019). For each module, the median number of genomes per family (m) was calculated. The genomes were ranked by the number of families they carry. The m genomes that carry the most of families were retained; their phyla distribution defines the taxonomic assignment of the module.

### Functional annotation

Protein sequences were functionally annotated based on the accession of their best Hmmsearch match (version 3.1) (E-value cut-off 0.001) (Eddy, 1998) against an HMM database constructed based on ortholog groups defined by the KEGG (Kanehisa et al., 2016) (downloaded on June 10, 2015). Domains were predicted using the same hmmsearch procedure against the Pfam database (version 31.0) (Punta et al., 2012). The domain architecture of each protein sequence was predicted using the DAMA software (version 1.0) (default parameters) (Bernardes et al., 2016). SIGNALP (version 5.0) (parameters: -format short-org arch) (Almagro Armenteros et al., 2019) was used to predict the putative cellular localization of the proteins. Prediction of transmembrane helices in proteins was performed using TMHMM (version 2.0) (default parameters) (Krogh et al., 2001). Protein sequences were also functionally annotated based on the accession of their best hmmsearch match (version 3.1) (E-value cut-off 1e-10) against the PDB database (Rose et al., 2017) (downloaded in February 2020).

### Sequence similarity analysis

The 75,737 subfamilies from the 10,866 families were searched against a bacterial database of 2,552 bacterial genomes (Méheust et al., 2019) using hmmsearch (version 3.1) (E-value cut-off 0.001) (Eddy, 1998). Among them 46,261 subfamilies, comprising 8,300 families, have at least one hit against a bacterial genome.

To calculate the ‘breadth’ metric, describing the incidence of archaeal protein families across domain Bacteria, we first found the percentage of representative genomes in each bacterial phylum with an above-threshold hit to each family. For those families with at least one hit among all bacterial reference genomes, we then secondarily calculated the percentage of unique bacterial phyla in which at least one third of representative genomes encoded a sequence hit. Finally, the breadth metric was compared with protein family size, taking into consideration KEGG categories derived from the functional annotation step described above, and used in part to identify systems of potential bacterial origin in DPANN (Figure 3 in main text).

## Data availability

The newly reconstructed genomes have been deposited at NCBI under BioProject **XX** (**To Be Announced**). The 569 genomes of the herein analysed archaea have been made publicly available on the ggkbase database (https://ggkbase.berkeley.edu/DPANN/organisms). Detailed annotations of the families and genomic contexts are provided in the Supplementary Dataset accompanying this paper. Raw data files (phylogenetic trees and fasta sequences) have been made available via figshare under the following link: https://doi.org/10.6084/m9.figshare.13517243.

## Author contributions

C.J.C., R.M. and J.F.B. designed the study. K.S., X.G. and B.J.B. collected the samples from the Guaymas Basin and assembled the genomes. C.J.C. and R.M. created the genome dataset. R.M. performed the protein family, the module detection, the genome annotations. C.J.C performed the phylogenetic and functional analyses. A.L.J. performed the bacterial analysis. C.J.C., R.M., A.L.J. and J.F.B. wrote the manuscript. All authors read and approved the final manuscript.

## Acknowledgments

We thank Dr. Christine He for the permission to use the metagenomic dataset from Genasci dairy farm.

