## Supplementary figures and images for "Protein family content uncovers lineage relationships and bacterial pathway maintenance mechanisms in DPANN archaea"

### FigureS1

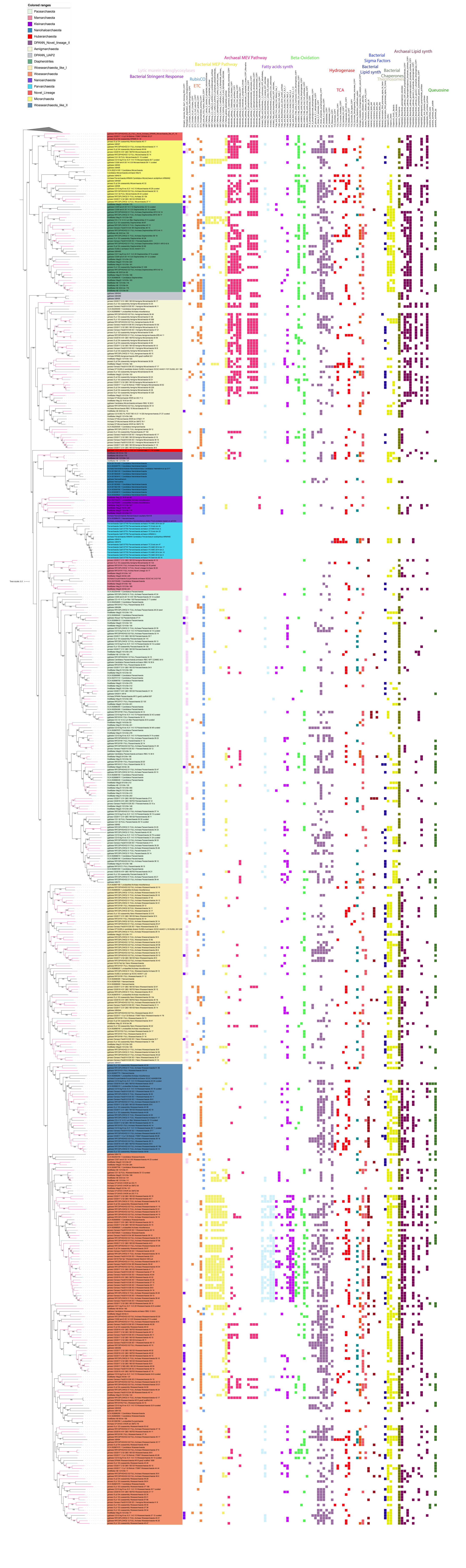
