## SupplementaryDataset_legend for "Protein family content uncovers lineage relationships and bacterial pathway maintenance mechanisms in DPANN archaea"

**Supplementary dataset**

**Table S1.** List of the 3,197 genomes used in this study. For each genome (column A), its NCBI accession, GGKBASE link, number of scaffolds, genome size and number of CDS are displayed in columns B, C, D, E and F respectively. Genome source is in column G, dRep cluster in column H. Genome completeness and the contamination based on single copy genes are displayed in columns I and J respectively. Column K informs about the concatenated ribosomal proteins. The 1,179 representative genomes are indicated in column L. The phylum and superphylum (DPANN and non-DPANN) taxonomy of the representative genomes are provided in columns M and N. Taxonomy based on the different databases we pulled out the genomes is shown in column O.

**Table S2.** Taxonomy distribution of the 17 modules. Module name is indicated in column A whereas the number of families is indicated in column B. Suggested taxonomic distribution is indicated in column C. Column D details the genomes used to define the taxonomic distribution (phylum, number of genomes).

**Table S3.** Annotation of the 10,866 families. Column A: module number. Column B: family accession. Column C: number of proteins in the family. Column D: median length of the proteins. Column E: ratio of proteins predicted to contain a signal peptide. Column F: median number of predicted transmembrane helix per protein. Column G: domain architecture reported by Pfam. Columns H, I, J, K, L: KEGG annotations. Column M: Cazy annotation. Columns N to AC indicate the ratio of genomes having the given family in the given archaeal phylum. Columns AD to CK indicate the ratio of genomes having the given family in the given bacterial phylum.

**Table S4.** Genes neighboring of the MEP pathway. The fifteen genes downstream and upstream of each fam03888 gene (column I) were identified and annotated using the protein clustering (column F), the PFAM (column H) and the KEGG databases (column G).

**Table S5.** Genes neighboring the three genes encoding the enzymes of the queuosine biosynthesis pathway. The seven genes downstream and upstream of each *QueA* (S-adenosylmethionine:tRNA ribosyltransferase-isomerase; fam24423), *QueH-like* (epoxyqueuosine reductase; fam24901) and *Tgt* (queuine tRNA-ribosyltransferase; Fam00366) gene (column H) were identified and annotated using the protein clustering (column E), the PFAM (column G) and the KEGG databases (column F).
